# A cotranslational role for selenocysteine in productive protein folding

**DOI:** 10.64898/2026.10.30.763106

**Authors:** Philip J. Robinson, Neil J. Bulleid, Kostas Tokatlidis

**Author notes:** Deceased (Neil J. Bulleid).

## Abstract

Selenocysteine (Sec) is generally regarded as the catalytic determinant that distinguishes selenoproteins from their cysteine (Cys)-containing counterparts. Consequently, the biological significance of Sec has largely been attributed to its unique chemical properties within mature proteins. Here, we uncover an unexpected role for Sec in protein biogenesis. Using the endoplasmic reticulum selenoprotein SELENOM as a model, we show that replacement of Sec with Cys redirects the protein into an alternative assembly pathway, resulting in formation of a stable domain-swapped homodimer. Biochemical and structural analyses demonstrate that this phenotype cannot be explained solely by altered active-site chemistry, as dimerisation persists following extensive active-site mutagenesis. Remarkably, reintroduction of Sec at alternative positions suppresses dimer formation, including when Sec is placed within a cleaved signal peptide and is absent from the mature protein. Thus, identical protein sequences can adopt distinct folding outcomes depending on whether Sec incorporation occurred during biosynthesis. These findings reveal a previously unrecognised role for Sec in productive protein folding and raise the possibility that evolutionary retention of the Sec incorporation pathway reflects advantages not only in catalysis but also in protein folding.

## Introduction

Selenoproteins contain the rare amino acid selenocysteine (Sec), which differs from the canonical amino acid cysteine (Cys) by the presence of selenium instead of sulfur in its side chain (Fig. 1A). There are only 25 selenoproteins in humans (Gladyshev *et al*, 2016) and these include the essential redox enzymes thioredoxin reductase 1 (Arner & Holmgren, 2000) and glutathione peroxidase 4 (GPX4) (Weaver & Skouta, 2022). Selenoproteins participate in a wide range of biological processes including antioxidant defence, redox signalling, endoplasmic reticulum (ER) homeostasis and calcium regulation (Ye *et al*, 2022). Because Sec possesses distinctive chemical properties, including enhanced nucleophilicity and a lower pKa than Cys (Johansson *et al*, 2005), studies of selenoproteins have traditionally focused on how Sec improves catalytic efficiency or alters enzymatic function. Consequently, the biological significance of Sec has largely been interpreted through the properties of the folded protein. By contrast, relatively little attention has been paid to whether the specialised pathway required to incorporate Sec might itself influence protein biogenesis. Incorporation of Sec occurs at what typically is a UGA stop codon (Donovan & Copeland, 2010). Recoding of this stop codon requires the presence of a downstream Sec insertion sequence (SECIS) element in the 3′ untranslated region of the mRNA (Copeland, 2003) (Fig.1B). The SECIS element recruits specialised elongation factors and a selenocysteinyl-tRNA, enabling co-translational reinterpretation of the UGA codon and insertion of Sec into the growing polypeptide chain (Labunskyy *et al*, 2014; Tico & Mariotti, 2026) (Fig. 1C).

**Figure 1.**
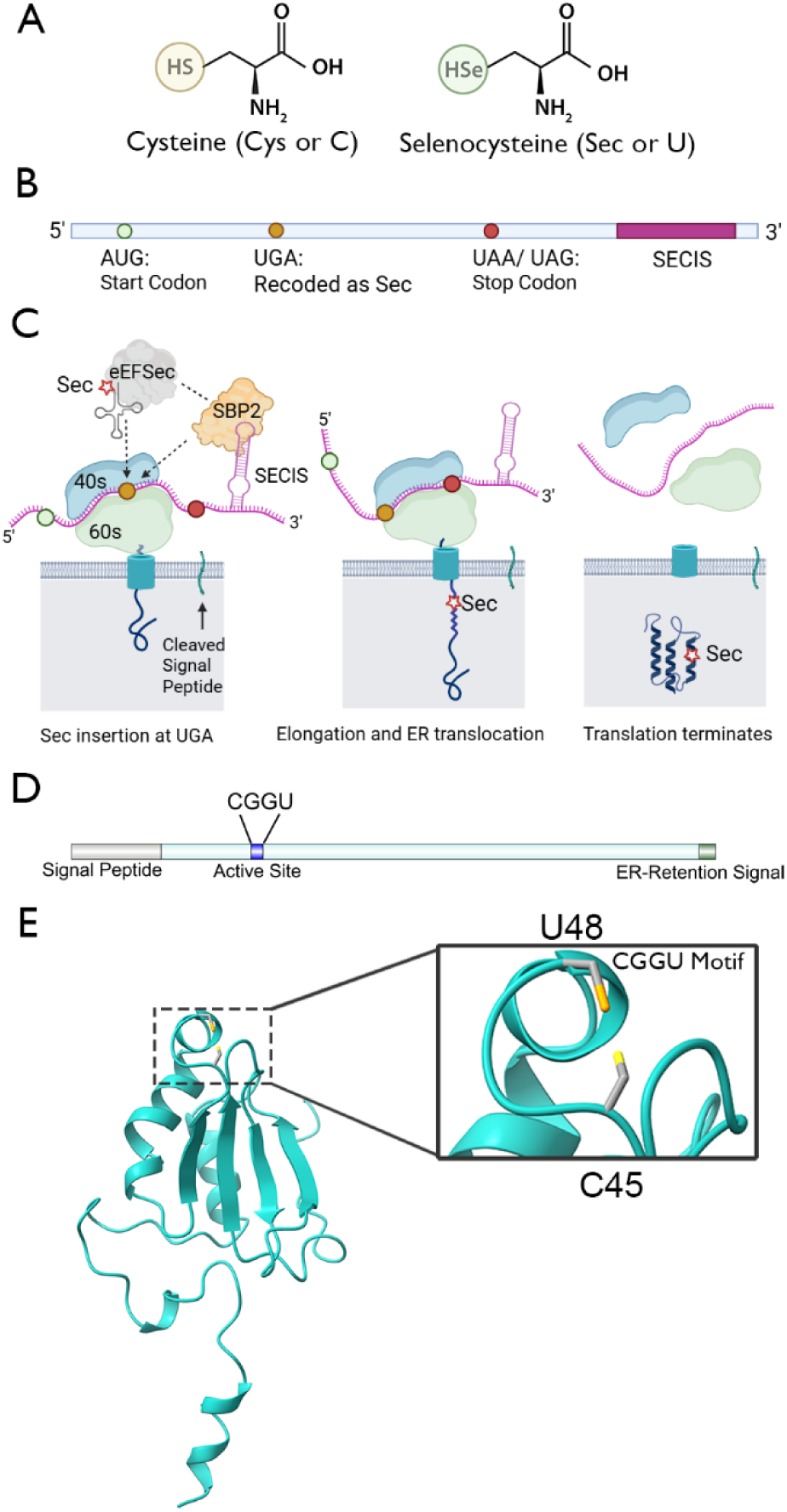
Selenocysteine incorporation mechanism and structural features of selenoprotein M (SELENOM). (A) Chemical structures of cysteine (Cys, C) and selenocysteine (Sec, U), highlighting the substitution of sulfur for selenium in the Sec side chain. (B) Schematic representation of a eukaryotic selenoprotein mRNA. Translation initiates at the AUG start codon, while an in-frame UGA codon is recoded to encode Sec. A SECIS (selenocysteine insertion sequence) element located within the 3′ untranslated region enables Sec incorporation, and translation terminates at a stop codon. (C) Mechanism of Sec incorporation during translation and translocation. The SECIS element is recognized by SECIS-binding protein 2 (SBP2), which recruits the Sec-specific elongation factor eEFSec and Sec-charged tRNA. This machinery promotes insertion of Sec at the UGA codon, allowing translation elongation and co-translational translocation of the nascent polypeptide into the endoplasmic reticulum (ER). Translation subsequently terminates at the stop codon, releasing the mature selenoprotein into the ER lumen. (D) Schematic representation of human selenoprotein M (SELENOM), showing the N-terminal signal peptide, the catalytic CGGU motif containing the Sec residue, and the C-terminal ER-retention signal. (E) Alphafold predicted structure of mature SELENOM following signal peptide cleavage, with a zoomed in view of the region containing the CGGU motif. The selenium atom is highlighted in orange and the sulfur atom in yellow. Panels A, B and C in this figure were drawn using Biorender.com

Sec incorporation is remarkably complex and inherently inefficient compared with the incorporation of canonical amino acids (Chavatte *et al*, 2025; Copeland & Howard, 2021). It requires dedicated RNA structures, specialised translation factors and recoding of a stop codon, and is associated with altered ribosome behaviour at UGA codons (Bohleber *et al*, 2022). Despite these costs, Sec has been repeatedly retained during evolution (Tico & Mariotti, 2026). The prevailing explanation is that selective pressure favours the superior redox chemistry of Sec relative to Cys (Arner, 2010). However, because Sec insertion modifies translational dynamics and nascent-chain synthesis, an alternative possibility is that the process of Sec incorporation contributes directly to protein folding and assembly. Whether specialised Sec incorporation influences productive folding pathways during protein biogenesis remains largely unexplored.

In humans, seven selenoproteins are located to the endoplasmic reticulum (ER). The ER is a specialised compartment for protein folding and the major site of disulfide bond formation within the cell (Ellgaard *et al*, 2016). Two of these ER-associated selenoproteins, SELENOF and SELENOM, are soluble luminal proteins that are structural homologues (Ferguson *et al*, 2006; Luo *et al*, 2026). Whereas SELENOF has an established role in glycoprotein quality control through its strong interaction with UGGT (Williams *et al*, 2024), the physiological function and molecular substrates of SELENOM remain poorly defined. Its redox-active CGGU motif, located within a thioredoxin-like fold (Fig. 1D,E), indicates a redox function (Ferguson *et al*., 2006; Luo *et al*., 2026). Although SELENOM is widely expressed, it is particularly abundant in the central nervous system suggesting an important role in neuronal physiology (Zhang & Song, 2021). It has been implicated in protection against oxidative stress and regulation of intracellular calcium homeostasis (Reeves *et al*, 2010) and has been shown to have redox activity (Gong *et al*, 2021). Recent studies have implicated SELENOM in ER proteostasis and responses to misfolded proteins, with SELENOM depletion activating ER stress pathways (Zhao *et al*, 2025). However, the molecular mechanisms underlying these effects remain unclear.

The involvement of SELENOM in ER proteostasis makes it an attractive model for investigating whether Sec incorporation influences folding trajectories during biosynthesis, beyond its established contribution to catalytic activity. Such effects may be especially apparent in ER proteins that operate within protein-quality-control pathways. Here, we identify proteins associated with SELENOM using complementary crosslinking and substrate-trapping proteomic approaches and investigate the structural consequences of Sec substitution. Unexpectedly, replacing Sec with Cys redirects SELENOM into an alternative assembly pathway consistent with formation of a stable domain-swapped homodimer. Strikingly, reintroduction of Sec at alternative locations, including within a signal peptide that is subsequently removed from the mature protein, suppresses dimer formation despite leaving the final protein sequence unchanged. These findings reveal an unanticipated role for Sec incorporation in directing productive folding of SELENOM and suggest that evolutionary selection for Sec may extend beyond catalytic function to include contributions to protein biogenesis.

## Results

### Selenium-dependent expression confirms efficient incorporation of Sec into recombinant SELENOM

To facilitate detection of SELENOM, a construct containing a C-terminal V5 epitope tag inserted immediately upstream of the ER-retention signal was generated (SELENOM-V5 (CGGU). SELENOM expression is known to be dependent on selenium availability (Chen *et al*, 2013; Sonet *et al*, 2021; Touat-Hamici *et al*, 2018). To assess this dependency, SELENOM-V5 (CGGU) was expressed in HT1080 cells cultured in the presence of increasing concentrations of sodium selenate, and protein expression was analysed by anti-V5 immunoblotting. Samples were analysed under reducing and non-reducing conditions to determine whether SELENOM formed stable redox-dependent complexes; however, no higher-molecular-weight species were observed. SELENOM-V5 (CGGU) was detected as a single species migrating at approximately 17 kDa (Fig. 2A). In the absence of added selenate, SELENOM-V5 (CGGU) expression was low, but increased markedly following selenate supplementation. Ǫuantification of the reduced protein revealed that expression increased approximately 8-fold at 1 μM sodium selenate and plateaued at approximately 12-15-fold above basal levels at 10-50 μM. These observations establish a system in which SELENOM expression responds to selenium availability as expected for a bona fide selenoprotein and provide the experimental framework for investigating how Sec influences SELENOM assembly. Based on these findings, sodium selenate was routinely included in growth media throughout the study, unless otherwise indicated, to ensure robust SELENOM expression.

**Figure 2.**
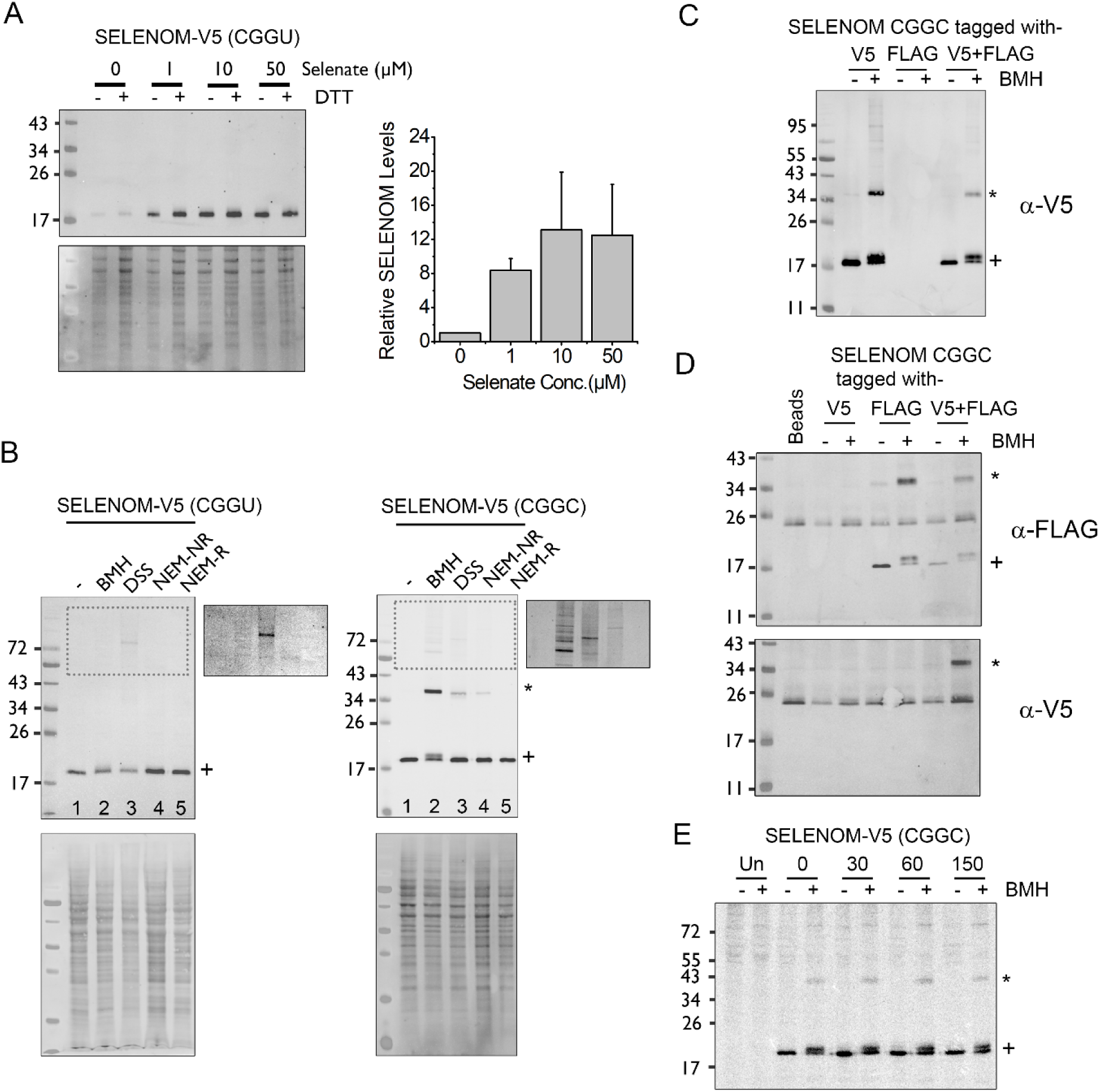
SELENOM overexpression and crosslinking analysis. (A) V5 immunoblot analysis of SELENOM-V5 (CGGU) expression in HT1080 cells grown in media supplemented with increasing concentrations of sodium selenate (0, 1, 10, or 50 μM). Lysates were analysed under non-reducing or reducing (+DTT) conditions as indicated. Total protein staining is shown as a loading control. SELENOM-V5 bands from the reducing samples were quantified (n=3) and plotted on the graph (right), error bars represent the standard deviation. (B) Assessment of SELENOM-V5 (CGGU) and SELENOM-V5 (CGGC) cellular interactions by chemical crosslinking and thiol-trapping approaches. Cells expressing the SELENOM-V5 variants were left untreated (−), treated with the crosslinkers BMH or DSS, or subjected to NEM treatment under non-reducing (NEM-NR) or reducing (NEM-R) gel conditions before V5 immunoblot analysis. The CGGC variant displays a crosslinked higher molecular weight species (asterisk) of approximately twice the monomer molecular weight, consistent with dimer formation. Monomeric SELENOM is indicated by (+). High contrast representations of the dotted box regions are shown to the right of each gel. Total protein staining is shown below as a loading control. (C) Cells expressing SELENOM (CGGC) tagged with either V5 or FLAG, or both together, were treated with or without BMH and cell lysates were analysed by reducing SDS-PAGE and immunoblotting using an anti-V5 antibody. The SELENOM monomer (+) and higher molecular weight species consistent with a dimer (*) are indicated. (D) Pull-down analysis of cells co-expressing V5- and FLAG-tagged SELENOM. Samples were treated with or without BMH, recovered on FLAG affinity beads, and analysed by immunoblotting with anti-FLAG (upper panel) and anti-V5 (lower panel) antibodies. The positions of monomeric SELENOM (+) and dimeric species (*) are indicated. (E) Pulse-chase analysis of SELENOM dimerisation. Cells expressing SELENOM were metabolically radiolabelled for 15 min, chased for the indicated times (0, 30, 60, or 150 min), treated with or without BMH, and analysed by SDS-PAGE followed by autoradiography. A crosslinked higher molecular weight species corresponding to a SELENOM dimer (asterisk) is detected throughout the chase period. An untransfected control (Un) is included. Monomeric SELENOM is indicated by (+).

### Substitution of SELENOM selenocysteine with cysteine promotes homo-dimer formation

To determine whether replacement of Sec affects SELENOM assembly, SELENOM-V5 (CGGU) and the Sec-to-Cys variant SELENOM-V5 (CGGC) were expressed in HT1080 cells and treated with the thiol-reactive crosslinker BMH or the amine-reactive crosslinker DSS. Lysates were analysed by immunoblotting to detect crosslinked species. Although substitution of Sec with Cys is generally considered conservative and is widely used to investigate selenoprotein function, it unexpectedly altered the oligomeric behaviour of SELENOM, suggesting entry into an alternative assembly pathway.

SELENOM-V5 (CGGU) remained predominantly monomeric following treatment with either BMH or DSS (Fig. 2B, lanes 1-3), with only a faint species migrating slightly above 72 kDa observed following DSS treatment. In contrast, SELENOM-V5 (CGGC) generated a prominent ∼34 kDa species following treatment with either crosslinker, consistent with dimer formation. Multiple additional higher-molecular-weight species were also detected above the 43 kDa marker (high contrast inset). The ∼34 kDa species was also observed in NEM-treated CGGC samples in the absence of crosslinker. This species was abolished under reducing conditions, indicating that the dimer is stabilised by an intermolecular disulfide bond.

The 34 kDa band present in the SELENOM-V5 (CGGC) samples is consistent with formation of a homodimer. To confirm this, SELENOM-V5 (CGGC) and SELENOM-FLAG (CGGC) were co-transfected into HT1080 cells, treated in the presence or absence of BMH, and isolated using anti-FLAG affinity beads. Samples were analysed by immunoblotting with anti-V5 and anti-FLAG antibodies and compared to controls expressing only a single tagged construct.

Expression of SELENOM-V5 (CGGC) was confirmed in whole-cell lysates by anti-V5 immunoblotting (Fig. 2C). A band corresponding to the SELENOM monomer was detected in all SELENOM-V5 (CGGC) expressing samples while a 34 kDa species consistent with a dimer band was evident exclusively after BMH treatment. Expression and isolation of SELENOM-FLAG (CGGC) following affinity purification was confirmed by anti-FLAG immunoblotting (Fig. 2D, upper panel).

To determine whether the 34 kDa species contained both tagged SELENOM protomers, FLAG affinity-purified samples were analysed using an anti-V5 antibody (Fig. 2D, lower panel). A 34 kDa V5-reactive band was detected only in samples co-expressing SELENOM-V5 (CGGC) and SELENOM-FLAG (CGGC) following BMH treatment. This species was absent from samples expressing either tagged construct alone. As anti-FLAG purification selectively isolates complexes containing SELENOM-FLAG, detection of this species with the anti-V5 antibody demonstrates that the complex contains both FLAG- and V5-tagged SELENOM molecules, confirming that the 34 kDa band corresponds to a SELENOM homodimer.

To assess the time scale over which dimer formation occurs, we performed a radiolabelled pulse chase assay using cells expressing SELENOM-V5 (CGGC) Fig. 2E. A crosslinked species corresponding to the SELENOM dimer was detected at the earliest chase time point examined (0 min, after a 15 min pulse) and remained detectable throughout the 150 min chase period (Fig. 2E). These findings indicate that dimer formation occurs rapidly following synthesis and are therefore consistent with an early folding or assembly event rather than progressive post-translational aggregation.

### Structural modelling suggests that Sec-to-Cys substitution redirects SELENOM into a domain-swapped assembly state

To investigate the structural basis of SELENOM-CGGC dimerization AlphaFold3 was used to model both monomeric and homodimeric forms of the protein. The monomer model adopts the thioredoxin-like fold expected for SELENOM, with the CGGC motif located within the loop connecting the first β-strand and first α-helix (Fig. 3A). The associated PAE map showed uniformly low predicted alignment errors across the structure, consistent with a high-confidence prediction (pTM = 0.81).

**Figure 3.**
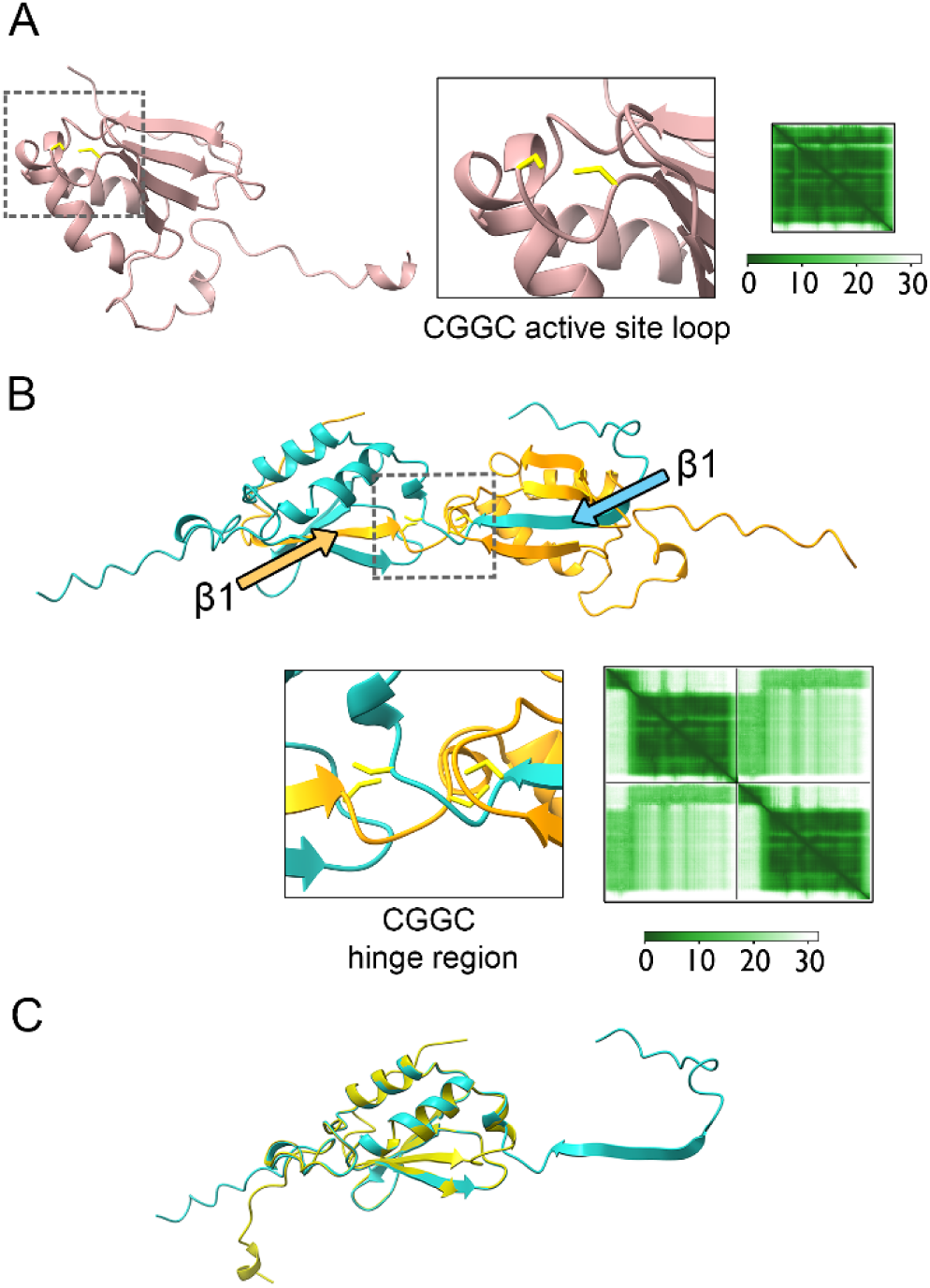
AlphaFold3 predicts a domain swap mechanism for the experimentally observed dimer. (A) AlphaFold3-predicted structure of monomeric SELENOM (CGGC). The overall fold is shown together with an enlarged view of the CGGC active-site loop (Cys in yellow) located within the thioredoxin-like domain. The accompanying predicted aligned error (PAE) map is shown on the right. (B) Representative AlphaFold3 multimer prediction of a SELENOM (CGGC) homodimer. Protomers are shown in cyan and orange. The first β-strand is exchanged between protomers(arrows), generating a domain-swapped interface centred on the CGGC hinge region. The boxed region is enlarged below, with Cys side chains shown in yellow. The corresponding PAE map is shown on the right. (C) Structural comparison of monomeric and dimeric SELENOM (CGGC) models. Overlay of the monomer model (green) with a representative dimeric protomer (cyan), showing that the overall fold is largely preserved and that structural differences are concentrated around the strand exchange region.

AlphaFold3 modelling of two protomers predicted a dimeric structure in which the β-strand immediately preceding the CGGC motif is exchanged between protomers, generating a domain-swapped dimer (Fig. 3B). Three of the five multimer predictions converged on a strand-exchanged architecture, while the remaining two models corresponded to non-interacting monomers (Fig. S2), suggesting that SELENOM-CGGC can adopt both monomeric and dimeric conformations, consistent with the mixed populations observed experimentally.

In the domain-swapped models, the CGGC motifs from each protomer are positioned within a hinge region at the dimer interface. The associated PAE map (Fig. 3B) showed relatively low predicted alignment errors within each protomer and a localised region of lower interchain error centred on the strand-exchanged interface, supporting a reproducible interchain interface despite uncertainty in the precise orientation of the protomers. The sulfur atoms of opposing Cys residues are separated by approximately 2.4 Å and 2.5 Å, distances compatible with intermolecular disulfide bond formation and within the range of BMH-mediated crosslinking observed in Fig.2. The model therefore provides a plausible structural explanation for the disulfide-linked homodimer observed experimentally.

To assess the structural consequences of dimerisation, a monomer model was superposed onto a protomer from the predicted dimer (Fig. 3C). The overall thioredoxin-like fold was largely preserved, with the exchanged β-strand occupying a similar position to that observed in the monomer structure. These observations suggest that dimerisation could occur through a local strand-exchange event without substantial rearrangement of the SELENOM core domain. Importantly, the model does not merely explain dimer formation but suggests that Sec-to-Cys substitution allows SELENOM to access this alternative conformational landscape that is normally avoided.

### Active-site chemistry alone cannot explain suppression of the dimeric state by Sec

The finding that substitution of Sec with Cys promotes SELENOM dimerization is surprising given that the two residues differ by only a single atom. One possible explanation is that replacement of Sec with Cys alters the redox properties of the active site and thereby favours dimer formation. To investigate this possibility, variants containing modifications of the CGGC motif were generated and analysed using the crosslinking and alkylation approaches (Fig. 4).

**Figure 4.**
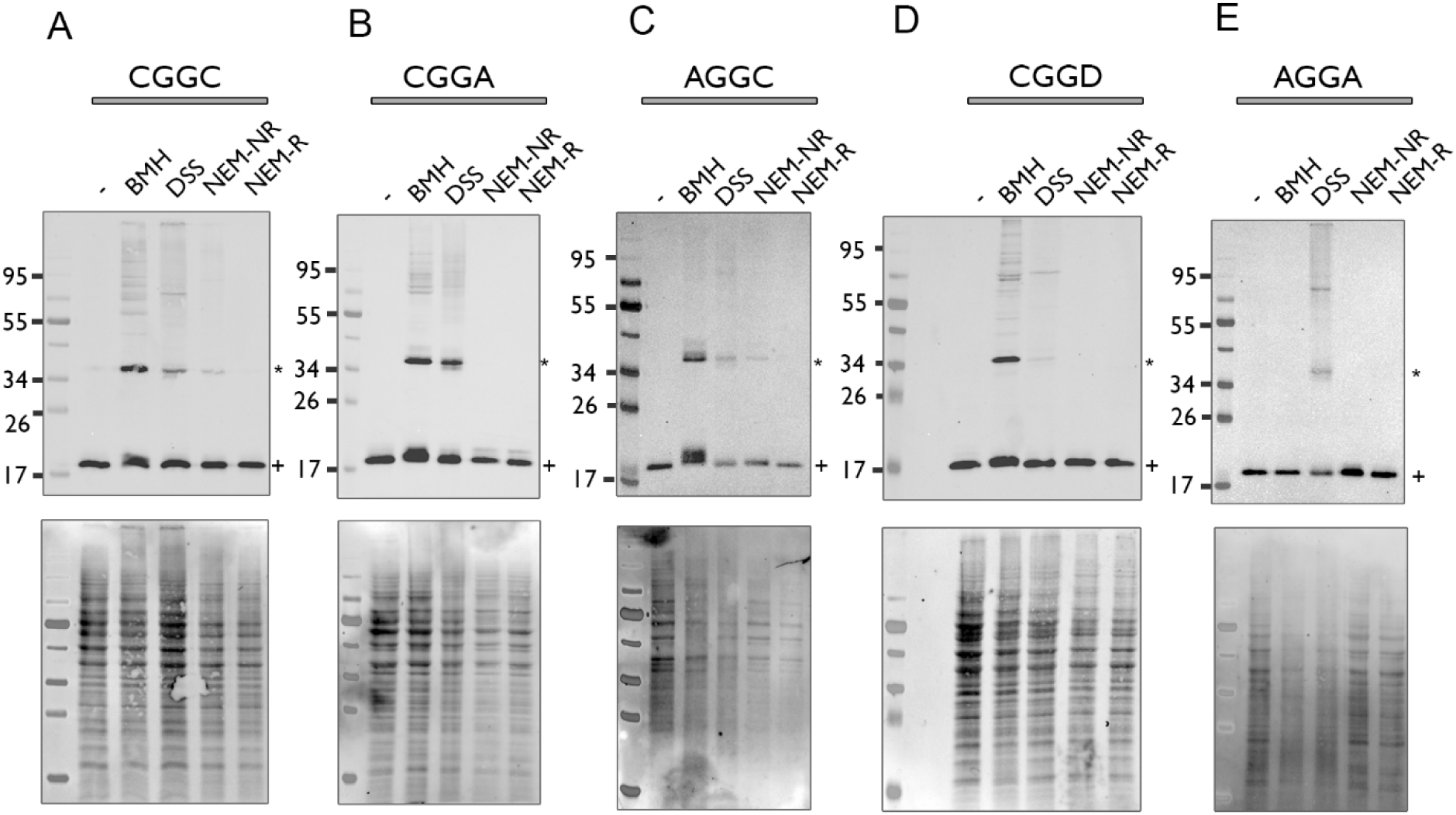
Dimerisation of SELENOM-V5 active-site mutants. V5 immunoblot analysis of SELENOM-V5 active site mutants: (A) CGGC, (B) CGGA, (C) AGGC, (D) CGGD, and (E) AGGA. Cells expressing V5-tagged SELENOM constructs were subjected to the indicated treatments prior to lysis: untreated (-), BMH, DSS and NEM. Samples treated with crosslinkers (BMH and DSS) were analysed by reducing SDS-PAGE, whereas NEM-treated samples were analysed under non-reducing (NR) or reducing (R) conditions. Immunoblotting with anti-V5 antibody detected SELENOM monomers migrating above 17 kDa (plus), together with dimeric species above ∼34 kDa (asterix) and higher molecular weight species observed under selected treatment conditions. Total protein staining shown below each blot serves as a loading control.

Mutation of either active-site Cys individually (CGGA or AGGC) did not prevent formation of either the ∼34 kDa dimer species or the higher molecular-weight crosslinked complexes (Fig. 4 B,C). In both cases, BMH treatment produced crosslinked species comparable to those observed for the CGGC mutant. These findings demonstrate that neither active-site Cys is individually required for dimer formation. The NEM-treated samples provided insight into the disulfide linkage responsible for the dimer. While the AGGC mutant retained a disulfide-linked dimer that was resolved under reducing conditions, the corresponding species was absent from the CGGA mutant. This suggests that the intermolecular disulfide bond is formed through the second Cys of the active-site motif.

Furthermore the lower pKa of Sec means it is likely to be negatively charged at cellular pH levels and this charge could inhibit dimerization. To investigate whether introduction of a negatively charged residue could suppress dimerization, a CGGD mutant was generated. This mutant continued to form a prominent dimer following BMH treatment (Fig. 4D) and displayed a crosslinking pattern similar to that observed for CGGA. Thus, introduction of a negatively charged residue at the active site is insufficient to prevent dimer formation.

Finally, an AGGA mutant lacking both active-site Cys residues was analysed. As expected for a protein lacking Cys and Sec, no BMH-dependent crosslinking or NEM-sensitive disulfide species were observed. Nevertheless, a dimeric species remained detectable following DSS treatment (Fig. 4E), indicating that dimer formation can occur in the complete absence of active-site thiols. It was noticed that the expression levels for the AGGA mutant were much lower (5%) than the other constructs (Fig. S3) and yet dimer formation is retained, arguing against a purely concentration dependent effect.

Collectively, these findings argue against a model in which dimerization arises solely from altered active-site redox chemistry. Instead, they suggest that the ability of Sec to suppress entry into the dimeric state reflects a mechanism distinct from its local catalytic properties.

### Selenocysteine directs productive folding independently of its position in SELENOM

The persistence of dimerisation following extensive active-site mutagenesis suggested that the effect of Sec may not simply reflect the redox properties of the active-site motif. We therefore investigated whether the unique mechanism of Sec incorporation itself might influence SELENOM folding. To test this, Sec residues were introduced into alternative positions within the SELENOM-V5 CGGA SECIS background (Fig. 5A). Two sites were selected: S35, located within the folded SELENOM domain but distant from the active site, and V18, located within the N-terminal signal peptide that is removed during ER translocation.

**Figure 5.**
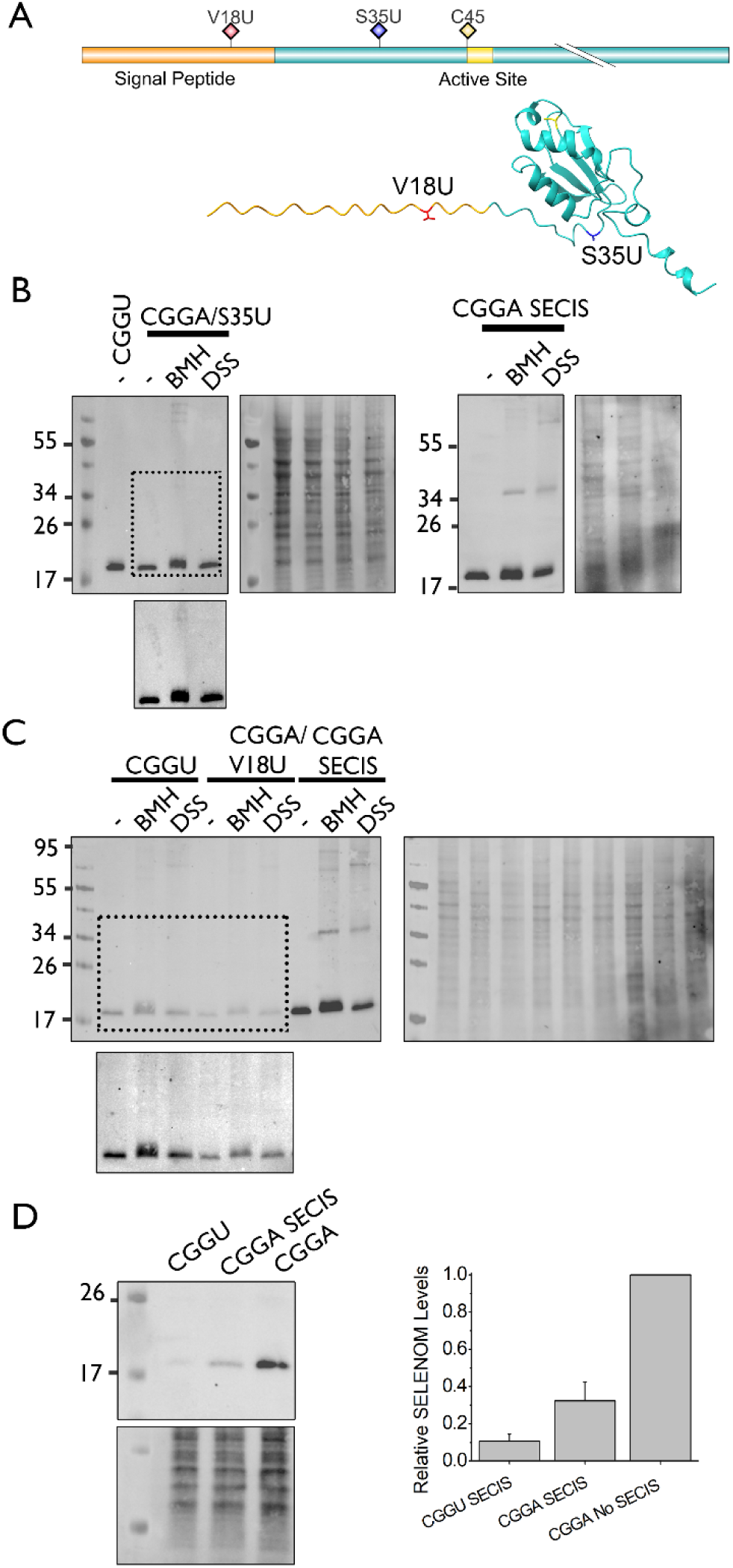
Position effects of selenocysteine on SELENOM dimerisation. (A) Schematic representation and structural model of SELENOM showing the signal peptide, active-site region, and positions of engineered amino acid substitutions. Selenocysteine (U) was introduced at residues V18 and S35 (V18U and S35U), while the native active-site cysteine (C45) is indicated. (B) Analysis of SELENOM S35U dimerisation. Cells expressing V5-tagged SELENOM CGGA S35U were treated with the crosslinkers BMH or DSS, alongside untreated controls. Expression was compared with wild-type SELENOM (CGGU). Crosslinking was compared with a Sec-deficient dimer control (CGGA SECIS). Left panels of each set show full anti-V5 immunoblots. Dotted boxes highlighting the region containing SELENOM monomer and dimers for the Sec containing proteins. The boxed region is shown at higher contrast. Right panels show total protein staining as a loading control. (C) Analysis of SELENOM V18U dimerisation. Cells expressing V5-tagged SELENOM CGGA V18U were treated with the crosslinkers BMH or DSS, alongside untreated controls. Mutant constructs were compared with wild-type SELENOM (CGGU) and with a Sec-deficient control (CGGA-SECIS). Left panel shows full anti-V5 immunoblots, with dotted boxes highlighting the region containing SELENOM monomer and dimers for the Sec containing proteins. The boxed region is shown at higher contrast. Right panel shows total protein staining as a loading control. (D) Effect of SECIS element inclusion on SELENOM expression. Anti-V5 immunoblot analysis of SELENOM constructs expressed from CGGU, CGGA+SECIS, or CGGA vectors. Total protein staining is shown as a loading control. The graph shows quantification of SELENOM expression from the different constructs. (n=3) error bars represent SD.

The CGGA/S35U construct was successfully expressed confirming incorporation of Sec (Fig. 5B). When treated with BMH or DSS, the S35U mutant showed little evidence of dimer formation. Although a faint higher molecular-weight species could be detected upon contrast enhancement, this signal was substantially reduced compared with the prominent dimer observed in CGGA-SECIS, which differs from S35U only by lacking the Sec incorporation site. Thus, suppression of dimerisation does not require Sec to occupy its native catalytic position, indicating that productive folding can be restored independently of active-site composition when Sec is introduced at an alternative position within the protein.

To further distinguish between effects of Sec chemistry and its insertion mechanism, Sec was introduced at position V18 within the signal peptide. Following signal-peptide cleavage, the mature V18U protein is predicted to be identical in sequence to the CGGA construct. Despite this, introduction of Sec at V18 almost completely suppressed dimer formation, yielding a migration pattern that resembled wild-type SELENOM rather than the dimerising CGGA-SECIS control (Fig. 5C).This result cannot readily be explained by altered catalytic activity, redox chemistry or structural contributions of Sec within the mature protein. Instead, it indicates that events occurring during Sec decoding and nascent-chain biogenesis influence the final folding outcome.

During these experiments, Sec-containing constructs consistently exhibited lower expression levels than their Cys-containing counterparts. In addition, the presence of the SECIS element appeared to reduce expression even in the absence of a UGA codon. To assess this directly, expression of CGGU, CGGA+SECIS and CGGA constructs was compared (Fig. 5D). Inclusion of the SECIS element lowered SELENOM expression relative to the CGGA construct, while introduction of a Sec codon caused a further decrease. These findings indicate that both the SECIS element and Sec incorporation influence overall expression efficiency. Notably, despite being expressed at only ∼5% of CGGA levels, the AGGA mutant still formed dimers (Fig. S3). Therefore, the failure of the Sec-containing variants to dimerise cannot be explained solely by reduced expression levels.

Together, these experiments demonstrate that Sec suppresses the off-pathway dimeric state independently of its location within mature SELENOM. Most strikingly, the ability of a transiently incorporated signal-peptide Sec residue to restore monomer formation indicates that the process of Sec incorporation itself contributes to productive folding. We therefore conclude that Sec functions not only as a catalytic residue but also as a determinant of productive protein folding, providing strong evidence that the biological significance of Sec can extend beyond its chemical contribution to the mature protein.

### SELENOM associates with ER pathways involved in oxidative folding and proteostasis

To identify proteins associated with SELENOM, the higher molecular weight species observed following BMH and DSS crosslinking of the CGGC variant (Fig. 2B) were isolated using anti-V5 affinity purification and analysed by mass spectrometry. To focus on candidate ER interaction partners, the dataset was filtered to retain proteins containing predicted signal peptides, and the relative abundances of proteins identified in both crosslinking datasets were compared (Fig. 6A).

**Figure 6.**
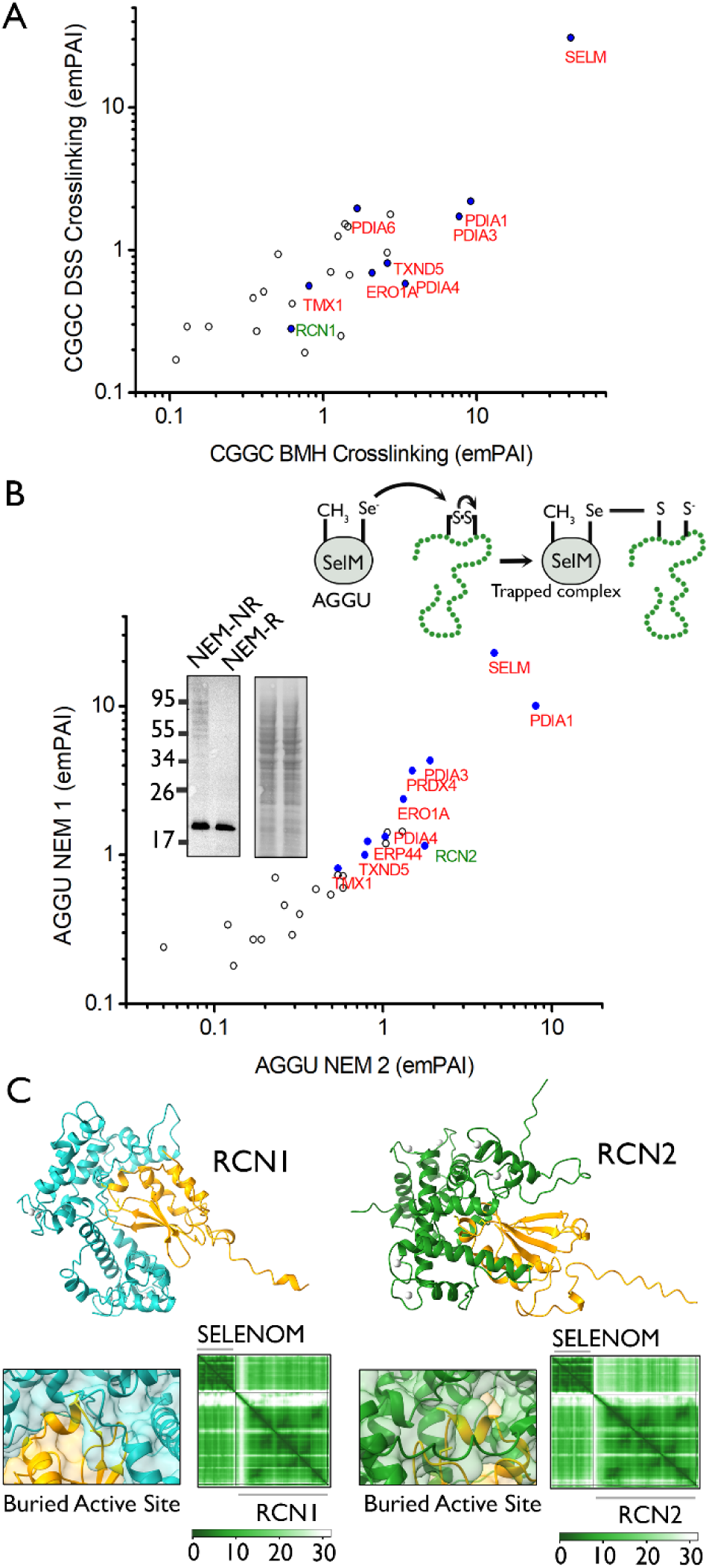
Identification of SELENOM-associated proteins by crosslinking and substrate-trapping proteomics and structural modelling of SELENOM-reticulocalbin interactions. **(A)** Comparison of proteins identified following crosslinking of wild-type SELENOM-V5 (CGGC) using BMH and DSS. Protein abundance is shown as emPAI values on logarithmic axes. ER oxidoreductases identified in both datasets are highlighted in red and RCN1 highlighted in green. **(B)** Comparison of proteins identified in two independent substrate-trapping experiments using the SELENOM-V5 AGGU mutant following NEM treatment. Protein abundance is shown as emPAI values on logarithmic axes. The diagram (top right) shows the mechanism of substrate trapping. The inset shows a representative V5-immunoblot (left) and total protein staining (right) demonstrating accumulation of higher molecular weight SELENOM-containing species under non-reducing gel conditions. Recurrent ER oxidoreductases identified in both experiments are highlighted in red and RCN2 highlighted in green. **(C)** AlphaFold3 predictions of SELENOM interactions with reticulocalbin-1 (RCN1) and reticulocalbin-2 (RCN2). Predicted complex structures are shown with SELENOM coloured orange and RCN proteins coloured blue (RCN1) or green (RCN2). Corresponding predicted aligned error (PAE) maps and zoomed in surface representations highlighting the buried active site are shown below each model.

Despite the distinct chemistries of the two crosslinkers, both datasets identified a similar group of ER oxidoreductases among the most abundant proteins recovered. These included PDIA1, PDIA3, PDIA4, PDIA6, ERO1A, TMX1 and TXND5 (Fig. 6A). The repeated identification of proteins involved in oxidative protein folding suggests that SELENOM associates with the ER redox network.

To complement the crosslinking experiments, we employed a substrate-trapping strategy using the SELENOM-V5 AGGU mutant. In this construct, replacement of the resolving Cys with Ala is predicted to stabilise mixed selenylsulfide intermediates formed during thiol-disulfide exchange reactions, enabling substrate capture (Fig. 6B). Following expression of this construct and treatment of cells with the alkylating agent NEM, non-reducing immunoblot analysis revealed SELENOM-containing higher molecular weight species that were resolved under reducing conditions (Fig. 6B, inset). Using anti-V5 affinity purification of NEM treated samples, two independent substrate-trapping experiments were compared and the hits identified in both samples plotted (Fig. 6B). This identified many of the same oxidoreductases observed in the crosslinking datasets, including PDIA1, PDIA3, PDIA4, ERO1A, TMX1 and TXNDC5. Additional redox-associated proteins, including PRDX4 and ERp44, were also detected. To assess the specificity of these interactions, a redox-inactive SELENOM-V5 (AGGA) control construct was analysed (Supplementary Excel File). Most of the recurrent oxidoreductases identified with AGGU, including PDIA1, PDIA4, TXNDC5 and ERO1A, were absent or greatly reduced in abundance. PDIA3 remained detectable but at substantially lower abundance than observed with the trapping mutant. These findings support the specificity of the interaction network identified by the AGGU substrate-trapping approach.

A further sample was tested expressing the wildtype protein (CGGU) with BMH crosslinker and was compared to the above data sets. Comparing the 5 data sets revealed a remarkable convergence of potential interaction partners (Table 1). The recurrence of the same proteins across independent crosslinking and substrate-trapping strategies increases confidence that these proteins represent biologically relevant SELENOM-associated factors. In particular, PDIA1, PDIA3, PDIA4, TXNDC5 and ERO1A were detected in all five experiments, including crosslinking analyses performed with both CGGC and wild-type SELENOM and the two independent substrate-trapping experiments. Several candidates, including PDIA1, PDIA3, CALR, CALX and GANAB, have also been identified previously in independent SELENOM interaction studies (Liu *et al*, 2024), further supporting their biological relevance. In addition to ER oxidoreductases, proteins associated with protein quality control, glycoprotein maturation and the calnexin cycle were repeatedly recovered. The enrichment of oxidative folding and quality-control factors is noteworthy given the observed effects of Sec on SELENOM assembly and supports a broader role for SELENOM within pathways that maintain protein-folding homeostasis

**Table 1.** Relative abundance of SELENOM and selected endoplasmic reticulum proteins identified in SELENOM-V5 immunoisolates under different crosslinking conditions. Rows highlighted in orange are proteins identified in all 5 samples. +indicates identification in a published crosslinking study (Liu *et al*., 2024).

|  | emPAI |  |  |  |  |  |
|---|---|---|---|---|---|---|
| Protein ID | CGGC DSS | CGGC BMH | CGGU BMH | AGGU NEM 1 | AGGU NEM 2 |  |
| SELM | 30.75 | 41.12 | 12.32 | 4.56 | 22.71 | Bait |
| RCN1+ | 0.28 | 0.62 | 0 | 1.34 | 0 | Calcium Homeostasis |
| RCN2+ | 0.9 | 0 | 0 | 1.77 | 1.15 |  |
| DJC10 | 0 | 1.07 | 0.11 | 0 | 0.17 | ER Oxidative Folding |
| ERO1A | 0.69 | 2.09 | 1.18 | 1.32 | 2.37 |  |
| PDIA1+ | 2.19 | 9.15 | 6.29 | 8.04 | 10.04 |  |
| PDIA3+ | 1.72 | 7.71 | 3.86 | 1.9 | 4.29 |  |
| PDIA4+ | 0.58 | 3.44 | 0.91 | 1.03 | 1.32 |  |
| PDIA6+ | 1.95 | 1.67 | 0 | 0 | 1.42 |  |
| TMX1 | 0.56 | 0.81 | 0 | 0.54 | 0.81 |  |
| TXND5 | 0.81 | 2.63 | 1 | 0.78 | 1 |  |
| CALR+ | 0 | 0.22 | 0.34 | 0 | 0.8 | Quality Control |
| CALX+ | 0.42 | 0.63 | 0.75 | 0 | 1.16 |  |
| ENPL | 1.52 | 1.39 | 1.78 | 1.26 | 0 |  |
| GANAB+ | 0.7 | 1.12 | 0.63 | 1.3 | 1.43 |  |
| HYOU1 | 0 | 0 | 0.14 | 0.29 | 0.29 |  |
| RPN1 | 1.46 | 1.45 | 0.74 | 0 | 0 |  |
| RPN2 | 0.51 | 0.41 | 0.31 | 0 | 0 |  |
| UGGG1+ | 0.17 | 0.11 | 0 | 0.17 | 0.27 |  |

Several proteins involved in extracellular matrix maturation, including LAMB3, PLOD2 and SERPINH1, were also identified in multiple datasets (Table S5). As these proteins are abundant secretory proteins in HT1080 cells, it remains unclear whether their recovery reflects direct SELENOM interactions or association with the broader ER protein-folding environment.

Among the other proteins identified in the screen the calcium binding CREC proteins reticulocalbin-1 (RCN1) and reticulocalbin-2 (RCN2) were notable because they were identified in multiple independent datasets and because SELENOM has previously been linked to calcium regulation. To explore these interactions further, AlphaFold3 modelling was performed. In contrast to most candidate interaction partners examined, AlphaFold3 consistently generated well-defined complexes between SELENOM and both RCN1 and RCN2 (Fig. 6C) with the models converging on similar interaction interfaces across independent runs. The predicted complexes positioned a similar surface of the SELENOM thioredoxin-like domain against each reticulocalbin and were associated with corresponding regions of low inter-chain PAE. Notably, the SELENOM active-site region became partially buried within the predicted interaction interface, raising the possibility that reticulocalbin binding could influence accessibility of the catalytic motif.

## Discussion

The principal finding of this study is that Sec influences the folding outcome of SELENOM independently of its established catalytic properties. Sec-to-Cys substitution redirects SELENOM into a stable alternative assembly state, whereas restoration of Sec, even at positions that do not contribute to the mature protein sequence, promotes productive monomer formation. This finding is unexpected because the Sec-to-Cys substitution is often regarded as a conservative change that preserves overall protein structure while modifying catalytic properties, and such mutants have been widely used to probe selenoprotein function (Johansson *et al*., 2005). Dimerisation persists following substitution of active-site Cys residues for Ala, arguing against a simple redox-related explanation. Most notably, introduction of Sec at alternative positions, including within a cleaved signal peptide, suppressed dimerisation despite leaving the mature protein sequence unchanged, suggesting that events associated with Sec incorporation influence the folding trajectory of SELENOM in the ER (Fig. 7).

**Figure 7.**
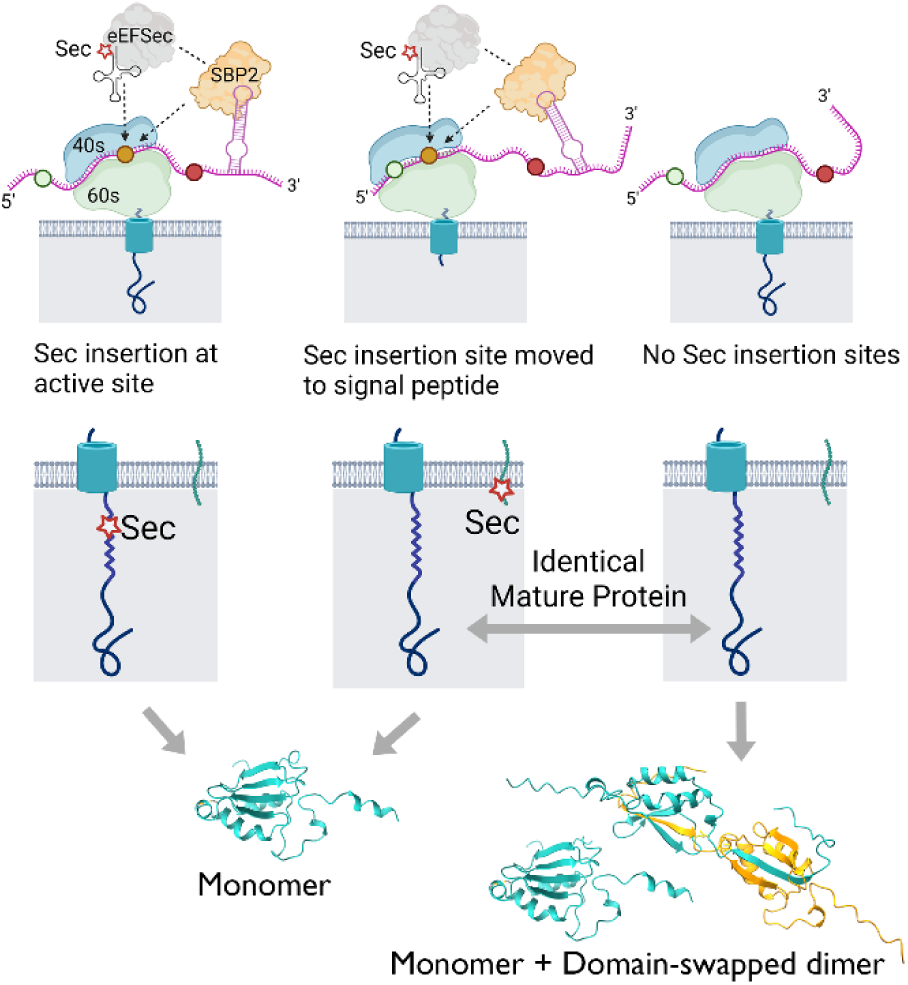
Model for the influence of selenocysteine incorporation on SELENOM folding. Selenocysteine (Sec) insertion recruits the SECIS-dependent translation machinery and yields mature SELENOM containing Sec within a monomeric thioredoxin-like fold. Relocation of the Sec codon to the signal peptide also favours formation of the monomeric state, despite the mature protein lacking Sec following signal peptide cleavage. In contrast, loss of Sec insertion through mutation promotes formation of a domain-swapped dimeric species. These findings suggest that the process of Sec incorporation during biosynthesis influences the subsequent folding and oligomerisation of SELENOM. Upper part of the figure was made using Biorender.com.

Modelling of the SELENOM dimer using AlphaFold3 produced a domain-swapped architecture involving β-strand exchange, which is a plausible model that fits with the experimental observations. Domain swapping is a well characterised structural phenomenon between homo-dimeric proteins (Lafita *et al*, 2019; Mascarenhas & Gosavi, 2017) and the same beta-strand swapping mechanism has been observed previously in a thioredoxin fold (Garcia-Pino *et al*, 2009). The model predicts that the active-site region contributes to the dimer interface, while biochemical analysis indicates formation of an intermolecular disulfide bond. Together, these observations suggest that dimerisation is likely to alter accessibility and reactivity of the catalytic motif relative to the monomeric state and imply that formation of the monomeric state may be functionally important.

We propose a model in which Sec incorporation biases SELENOM toward a productive folding trajectory during biogenesis. Under this model, the specialised and highly regulated machinery required for UGA recoding (Peng *et al*, 2021) transiently alters translation dynamics (Bohleber *et al*., 2022; Copeland & Howard, 2021), and thereby influences the conformational landscape available to the nascent polypeptide. Protein folding trajectories are highly sensitive to translation kinetics, and numerous studies have demonstrated that synonymous mutations can alter protein folding by modifying cotranslational synthesis rates (Liu *et al*, 2021). In the absence of Sec incorporation, SELENOM more readily enters an alternative pathway culminating in formation of a domain-swapped dimer. Although the molecular basis of this effect remains to be established, the V18U experiment demonstrates that the phenomenon originates during biogenesis rather than from chemical properties of the mature protein itself.

These observations suggest that the specialised translational pathway required for Sec incorporation can influence the structural fate of the nascent polypeptide.

An additional observation was that incorporation of the SECIS element reduced SELENOM expression even in the absence of a UGA codon, while introduction of a Sec codon caused a further decrease. Sec insertion is known to be inefficient (Vindry *et al*, 2018) and wild-type SELENOM (CGGU) was expressed at approximately 10% of the level observed for the CGGA mutant. Although the basis of these effects remains unclear, it indicates that both the SECIS element and the recoding process influence expression. Reduced expression may influence intracellular protein abundance and therefore affect the relative populations of monomeric and oligomeric species (Gell *et al*, 2012). However, the AGGA mutant still forms dimers despite being expressed at substantially lower levels than the Sec-containing variants, suggesting that altered expression alone cannot account for the observed differences in dimerisation. Instead our results favour a model where altered folding trajectories due to the unusual Sec insertion mechanism result in an altered oligomeric state (Fig.7).

Several recurrent SELENOM-associated proteins identified in the present study, including PDI family members, calnexin and calreticulin and the reticulocalbins, were also reported in an independent photo-crosslinking study (Liu *et al*., 2024). These overlapping findings support an association between SELENOM and ER protein folding pathways. In contrast, the repeated identification of ERO1A, TXND5, TMX1 and DNAJC10 in our datasets expands the known SELENOM interaction network further. PDI family members are known to form a disulfide exchange network (Fass & Thorpe, 2018; Oka *et al*, 2015) and further studies are needed to elucidate SELENOM’s function in these pathways. The simultaneous identification of oxidoreductases and glycoprotein quality-control factors is notable because these pathways are functionally interconnected (Ellgaard *et al*., 2016), raising the possibility that SELENOM participates at the interface between these pathways. The SELENOM homologue SELENOF, which functions in association with the UGGT complex (Williams *et al*., 2024), may suggest a similar specialised role for SELENOM in ER quality-control pathways.

Our previous work has characterised the reductive pathway in the ER that is required for reducing and rearranging disulfide bonds in folding proteins (Cao *et al*, 2020; Poet *et al*, 2017). Two of the oxidoreductase enzymes identified here, ERdj5 (DNAJC10) and Erp57 (PDIA3), reduce non-native disulfide bonds (Oka *et al*, 2013; Robinson *et al*, 2023) but the upstream reductase systems responsible for maintaining these enzymes in their active reduced state remains unclear. As Sec is typically an excellent nucleophile (Arner, 2010) and as SELENOM’s structure suggests it is a redox enzyme (Ferguson *et al*., 2006), it represents an attractive candidate for this purpose. The proteomics data with the substrate trap mutant support this, as the nature of the mechanism traps substrates undergoing reduction (Jessop *et al*, 2009), and so the identification of multiple PDI family members through this process is consistent with SELENOM functioning within this reductive network and raises the possibility that SELENOM acts as an upstream reductase.

Given previous reports linking SELENOM to calcium homeostasis (Liu *et al*., 2024; Reeves *et al*., 2010), and the established role of SELENON as a reductase and calcium sensor (Chernorudskiy *et al*, 2020) the identification of the calcium-binding ER luminal CREC family proteins (Honore, 2009) RCN1 and RCN2 is particularly intriguing. Although their functional relationship with SELENOM remains unclear, the reproducible AlphaFold3 interaction models and predicted burial of the SELENOM active site suggest a potential regulatory interaction that warrants further investigation.

Collectively, our findings reveal an unexpected function for Sec in protein biogenesis. Beyond providing distinctive catalytic chemistry, Sec can influence the structural fate of a protein during its synthesis. The observation that a transiently incorporated Sec residue within a cleaved signal peptide is sufficient to restore productive folding suggests that the process of Sec incorporation itself carries biological information. These findings raise the possibility that evolutionary retention of the complex Sec incorporation machinery reflects advantages in protein folding and structural maturation in addition to the well-established catalytic benefits (Arner, 2010) of the twenty-first amino acid.

## Materials and Methods

### Plasmid Design and Site Directed Mutagenesis

A plasmid was designed encoding the wildtype human SELENOM sequence (Uniprot: Ǫ8WWX9) with a V5 tag before the C-terminal ER retention signal. The SELENOM 3′ untranslated region containing the endogenous SECIS element was retained to permit Sec incorporation. This plasmid was called SELENOM-V5 (CGGU). As multiple variants of the SELENOM CGGU active-site motif are examined throughout this study, the composition of this motif is always indicated. A Cys-substituted variant, SELENOM-V5 (CGGC), was generated by replacement of the active-site Sec residue and removal of the SECIS-containing 3′ untranslated region. A FLAG-tagged version of the SELENOM CGGC construct, designated SELENOM-FLAG (CGGC), was also generated. These constructs were synthesized as inserts in pcDNA3.1 plasmid using GeneArtTM gene synthesis (Thermo Fisher Scientific), with the sequence of each insert provided in Fig S1. Using the above plasmids as templates, mutants were generated (Table S1) by site-directed mutagenesis (SDM) using ACCUZYME^TM^ (Bioline), with the details of the primers and templates used for this purpose in Table S2.

### Cell Culture

HT1080 (CCL-121) human fibrosarcoma cells were purchased from the American Type Culture Collection (ATCC) in 2022 (authenticated by the supplier). A cell bank was made on receipt of this stock with aliquots taken as needed. Cells were cultured in DMEM (Gibco) supplemented with 2 mM glutamine (Gibco), 100 U/ml penicillin (Sigma), 100 µg/ml streptomycin (Sigma) and 10 % foetal calf serum (Sigma) at 37 °C in a 5 % CO2 incubator. Cells in culture were regularly tested for mycoplasma contamination.

### Transfection

HT1080 cells at ∼80% confluence were transfected using MegaTran 2.0 transfection reagent (OriGene, Cambridge, UK). For this purpose, MegaTran was mixed with plasmid DNA in a 3:1 ratio (µl reagent/ µg DNA) in serum-free DMEM, before dropwise addition to the cultured cell media. After ∼16 hours, the medium was replaced. For cells expressing SELENOM variants containing Sec and their associated controls (Experiments show in Fig 2B, Fig 5. B-D, and all proteomics samples) 50 µM sodium selenate was also added at this stage. Cells were then incubated for a further 3 hours before processing. For SELENOM expression analysis (Fig. 2A), a modified protocol was used in which transfection reagents were removed after 8 h and replaced with fresh medium containing the appropriate concentration of sodium selenate, followed by overnight incubation prior to further analysis.

### Crosslinking and NEM treatment

For crosslinking reactions, transfected cells were trypsinised, treated with Soybean Trypsin Inhibitor (Sigma) at 100 µg/ml for 30 s on ice and isolated by centrifugation for 1 min at 1,000 x g. Cell pellets were washed with PBS and resuspended in 20 µl of 20 mM HEPES (pH 7.4) before treatment with 1 mM BMH or DSS crosslinker (prepared from 20 mM stocks in DMSO). Reactions were incubated on ice for 10 min and quenched with 10 mM DTT (BMH reactions) or 100 mM Tris-HCL (pH 8.0) (DSS reactions). Cells were then pelleted (1 min at 1,000 x g) and washed with 20 mM HEPES (pH 7.4). The cell pellets were resuspended in lysis buffer ((50 mM Tris-HCl pH 7.5, 1% (v/v) Triton X-100, 150 mM NaCl, 2 mM EDTA, 0.5 mM PMSF, and 0.02 % (w/v) sodium azide, with an EDTA free protease inhibitor tablet added) and incubated on ice for 15 mins, before centrifugation at 16,000 g for 10 mins. The resulting supernatants were used for further analysis.

For samples in which disulfide-linked species were analysed, transfected cells were washed with PBS and treated with 25 mM NEM in PBS for 10 min on ice. Lysis buffer was then added directly to the cells, which were harvested by scraping and incubated on ice for 15 min before centrifugation at 16,000 × g for 10 min. The resulting supernatants were used for further analysis.

### Immunoisolation Analysis

SELENOM-V5 (CGGC) and SELENOM-FLAG (CGGC) were transfected individually or co-transfected into cells and crosslinked with BMH as described above. Following cell lysis the supernatant was taken, 10 % of which was retained for immunoblot analysis and the remainder was precleared (30 min at 4 °C) using protein A Sepharose (PAS) (1 % v/v) (Generon, UK). Following centrifugation, the supernatant was taken and incubated with 20 µl of Anti-FLAG M2 affinity Agarose (Cat# A2220, Sigma) for ∼16 hours. Samples were washed three times with lysis buffer, and once with 20 mM HEPES, pH 7.4.

### SDS-PAGE and Immunoblotting

Supernatants from cell lysates or immunoisolated complexes were diluted in reducing/non reducing SDS-PAGE sample buffer and boiled (5 min). Samples were cooled and treated with NEM (50 mM). Samples were resolved by 15 % SDS-PAGE and proteins were transferred to a nitrocellulose membrane (0.45 µm) (GE Healthcare) via a wet transfer. The total protein transferred to membranes was assessed using TotalStain Ǫ (Azure Biosystems, cat#AC2227) according to the manufacturer’s instructions. The membranes were then blocked in 3 % w/v dried skimmed milk (Milfresh) in TBST buffer (50 mM Tris pH 7.6, 150 mM NaCl, 0.1 % Tween) for 1 hour. The blocked membranes were incubated with anti-V5 antibody (Invitrogen, cat#R960-25; 1: 500 dilution) or anti-FLAG-M2 antibody (Merck/Sigma) (cat #F3165, 1:500 dilution) for 16 hours. After primary antibody incubation, the membranes were washed (3 x, 5 min) with TBST and incubated with goat anti-mouse DyLight™ 800-conjugated secondary antibody (Invitrogen, cat #SA535521; 1:10,000 dilution) for 30 min at room temperature. The membranes were washed (3 x, 5 min) with TBST and were scanned using an Azure c600 imaging system. Ǫuantification was performed using ImageJ 1.53K (NIH), with signal normalised to total protein. Statistical tests are indicated in the corresponding figure legends. All immunoblot-based experiments were repeated at least 3 times unless otherwise indicated and representative blots are shown.

### Proteomic Analysis by LC-MS/MS

Following crosslinking or NEM treatment protocols described above, and subsequent cell lysis, the supernatant was taken and pre-cleared using PAS (1 % v/v). The samples were centrifuged, and the resulting supernatant was incubated with approximately 15 µl Anti-V5 Agarose Affinity Gel (Sigma, cat# A7345) overnight. Samples were prepared for proteomics by washing the beads with 3x lysis buffer, 3x lysis buffer but with 0.1% triton and 5x lysis buffer but without triton.

SELENOM-V5 complexes bound to V5 affinity agarose beads were sent to BSRC Mass Spectrometry and proteomics Facility, St Andrews UK, for analysis. The beads were resuspended in 20 mM Ammonium Bicarbonate, and subjected to trypsin digestion (Sigma-Aldrich, Cat. No. T4799), at 30°C overnight. The supernatant was recovered and concentrated to 20 µl using a Speedvac.

Peptides were subjected to LCMS/MS using an Ultimate 3000 RSLC (Thermo Fisher Scientific) coupled to an Orbitrap Fusion Lumos mass spectrometer with a FAIMS interface (Thermo Fisher Scientific). Peptides were injected onto a reverse-phase trap (Pepmap100 C18 5 μm,100 Å, 100 µm × 2cm) for pre-concentration and desalted with loading buffer (100% water, 0.05% TFA), at 5 μL/min for 10 min. The peptide trap was then switched into line with the analytical column (Easy-spray Pepmap RSLC C18 2 μm, 15 cm x 75 μm ID). Peptides were separated using mobile phase A (0.1 % formic acid in water) and mobile phase B (0.1% formic acid in 80% acetonitrile). Peptides were eluted using a linear gradient of 2-40 % B over 40 min, followed by an increase to 95 % B over 4 min. The column was washed at 95 % B for 6 min and subsequently re-equilibrated at 2% B for 10 min. The FAIMS interface was set to -45V and -65V at standard resolution. The mass spectrometer was operated in DDA positive ion mode with a cycle time of 1.5 s. The Orbitrap was selected as the MS1 detector at a resolution of 120,000 with a scan range of from m/z 375 to 1500. Peptides with charge states 2 to 6 were selected for fragmentation in the Iontrap using HCD as collision energy.

The raw data files were converted into mgf using MSconvert (ProteoWizard) and searched using Mascot 2.8.1 with trypsin specified as the cleavage enzyme and one missed cleavage permitted. Oxidation of methionine was set as a variable modification. For NEM treated samples, NEM modification of Cys residues was included as a fixed or variable search modification. Searches were performed against the SwissProt_2020_04 database. The precursor and fragment ion mass tolerances were set to 20 ppm and 0.6 Da, respectively.

Proteins identified by Mascot were retained if they achieved a protein score greater than 50 and were supported by at least two unique peptide sequences. emPAI values were compared with those obtained from an un-transfected control sample, and proteins enriched by at least two-fold were retained for further analysis. Protein identifiers were mapped using UniProt, and proteins containing a predicted signal peptide were selected to enrich for proteins entering the endoplasmic reticulum (ER). The resulting protein sets were visualised according to their emPAI values, with complete datasets provided as a supplementary file. Table 1 was generated from proteins detected in three or more independent datasets and were subsequently grouped according to their annotated biological functions.

### Pulse chase labelling

Cells were cultured in 6 well plates and were transfected with SELENOM-V5 CGGC overnight. Untransfected cells were cultured in parallel for comparison. The following day the media was replaced to remove the transfection reagents and the cells were incubated for a further 3 hours. To initiate the starvation phase, cells were incubated in Met- and Cys-free DMEM for 15 min. The same media supplemented with 11 μCi/ml of EXPRESS 35S Protein Labelling Mix (PerkinElmer Life Sciences) was then added and incubated for 15 mins to initiate the pulse phase. Following the pulse, the radiolabel-containing medium was removed, cells were washed with PBS, and complete medium was added to initiate the chase phase.

At each time point (0, 30, 60 or 150 min), samples were washed with PBS and trypsinised, before treatment with STI and crosslinking with BMH as described above. Samples were then centrifuged, and resuspended and incubated in lysis buffer (15 min on ice). Following centrifugation (16,000 × g for 10 min), supernatants were pre-cleared with PAS (1% v/v) for 30 min. Samples were then incubated overnight at 4°C with rabbit anti-V5 serum (kindly provided by Stephen High, University of Manchester; 1:100 dilution) and PAS (1% v/v) for immunoisolation. Immunocomplexes were washed 3 times with lysis buffer, and once with 20 mM HEPES (pH 7.4) and resuspended in reducing sample buffer prior to boiling. Samples were resolved by SDS-PAGE and gels were fixed in 10 % (v/v) TCA/10 % (v/v) methanol, dried and exposed to a phosphorimager plate and analysed on a FLA-7000 bioimager (Fujifilm). The experiment was repeated 3 times independently and a representative gel is shown.

### AlphaFold3 Modelling

Structures of SELENOM monomers, SELENOM dimers and SELENOM in complex with reticulocalbins were modelled using AlphaFold3 (Abramson *et al*, 2024). This was performed using the mature protein sequences following removal of the annotated N-terminal signal peptides from UniProt entries Ǫ8WWX9 (SELENOM), Ǫ15293 (RCN1), and Ǫ14257 (RCN2).For the reticulocalbin complexes, six calcium ions were included in each model. In all cases five models were generated. Confidence metrics including pTM, ipTM, and predicted aligned error (PAE) were used to assess prediction reliability (Tables S3 and S4). Structural visualizations of SELENOM containing Sec in Figures 1, 5 and 7 were generated using the published model deposited in ModelArchive under accession DOI: <u>10.5452/ma-j0za1</u>

### Declaration of generative AI and AI-assisted technologies in the writing process

Microsoft 365 Copilot was used during the writing of this article to improve the quality of the text. The outputs were scrutinised and edited as necessary. The authors take full responsibility for the content of the article.

### Software

Bar charts were prepared using Origin 6.0 professional software. Molecular graphics images were drawn with UCSF ChimeraX version 1.9 (Meng *et al*, 2023). *IBS 2.0 (Xie et al, 2022) was used to draw protein diagrams in Fig 1D and Fig. 5A.* Figure panels that were made in Biorender are available from the links below.

*Robinson, P. (202C)* https://BioRender.com/1kkyf7t, *Robinson, P. (202C)* https://BioRender.com/0ig8tel *Robinson, P. (202C)* https://BioRender.com/Sbx4ne5

## Acknowledgements

The authors wish to acknowledge Richard Cogdell, Brian Smith and Bill Cushley, and Martin Rennie (University of Glasgow) for advice throughout the project. Current and former members of the Bulleid group especially Marcel van Lith and Ojore Oka for discussion on the project and Bethany Fleming and Claudia Chieruzzi for technical support. We thank the BSRC Mass Spectrometry and Proteomics Facility, University of St Andrews, UK for analysis of samples and advice with sample preparation. The instrument used was funded by the BBSRC BB/T107686/1. This project was funded by a Wellcome Discovery Award 220253/Z/20/Z awarded to Neil Bulleid.

